# Nucleus accumbens volume is related to obesity measures in an age-dependent fashion

**DOI:** 10.1101/773119

**Authors:** Isabel García-García, Filip Morys, Alain Dagher

## Abstract

Motivation theories of obesity suggest that one of the brain mechanisms underlying pathological eating and weight gain is the dysregulation of dopaminergic circuits. While these dysregulations occur likely at the microscopic level, studies on gray matter volume reported macroscopic differences associated with obesity. One region suggested to play a key role in the pathophysiology of obesity is the nucleus accumbens (NAcc). We performed a meta-analysis of findings regarding NAcc volume and overweight/obesity. We additionally examined whether gray matter volume in the NAcc and other mesolimbic areas depends on the longitudinal trajectory of obesity, using the UK Biobank dataset. To this end, we analysed the data using a latent growth model, which identifies whether certain variables of interest (e.g. NAcc volume) is related to another variable’s (BMI) initial values or longitudinal trajectories. Our meta-analysis showed that, overall, NAcc volume is positively related to BMI. However, further analyses revealed that the relationship between NAcc volume and BMI is dependent on age. For younger individuals such relationship is positive, while for older adults it is negative. This was corroborated by our analysis in the UK Biobank dataset, which includes older adults, where we found that higher BMI was associated with lower NAcc and thalamus volume. Overall, our study suggests that increased NAcc volume in young age might be a vulnerability factor for obesity, while in the older age decreased NAcc volume with increased BMI might be an effect of prolonged influences of neuroinflammation on the brain.

## Introduction

Obesity may result in part from an increased appetitive drive interacting with an obesogenic environment (Michaud et al., 2017). It has been suggested that a trait, uncontrolled eating, may result from the interaction of increased reward drive and reduced self-control. Neural correlates of the uncontrolled eating trait may be found in mesolimbic circuits in the brain (Vainik et al., 2019).

The nucleus accumbens (NAcc) is a key node of the mesolimbic circuits (Haber and Behrens, 2014). The activation of these circuits signals rewards and sustains goal-directed behavior (Grace et al., 2007; Haber and Knutson, 2010). Neuropsychiatric studies have suggested that altered functionality in these circuits can give rise to impulsive and compulsive symptoms (Brooks et al., 2017). In the same vein, some types of obesity could theoretically stem from dysfunctions in the NAcc and other mesolimbic areas (Volkow et al., 2013).

Several studies have examined the relationship between adiposity and gray matter volume in the NAcc and other mesolimbic areas. Results, however, are heterogeneous. Some works have found *positive* associations between BMI and NAcc volume (Coveleskie et al., 2015; Horstmann et al., 2011). By contrast, a recent study using the UK Biobank cohort has reported that, in men, percentage of body fat is *negatively* associated with the volume of the NAcc along with other reward-related regions (e.g., caudate, pallidum, thalamus and hippocampus) (Dekkers et al., 2019). In women, only the pallidum shows negative associations with body fat (Dekkers et al., 2019). Null findings in the relationship between NAcc volume and BMI have also been reported. In fact, a coordinate-based meta-analysis on adiposity and gray matter volume in the whole brain did not find any differences in the NAcc (García-García et al., 2018). In the meta-analysis, the medial prefrontal cortex (extending to OFC) was the only reward-related region that showed (negative) correlations with adiposity (García-García et al., 2018).

Conflicting findings might be related to two different mechanisms underlying the relationship between NAcc volume and excess weight. First, increased NAcc volume might be a vulnerability factor for the development of obesity. In fact, it has been shown that striatal activity in response to foods seems to predict longitudinal weight gain (Demos et al., 2012; Geha et al., 2013; Stice et al., 2008; but see Stice and Yokum, 2018). Although the specific mechanism linking NAcc volume to reward function is unknown, it is possible that increased NAcc volume reflects enhanced appetitive drive. Second, in addition to being a vulnerability factor for obesity, NAcc volume might also be affected by chronic excess weight. Overall, overweight and obesity are associated with metabolic and cardiovascular pathology (Bastien et al., 2014) and are related to grey matter atrophy (Raji et al., 2009). The accumulation of adiposity can trigger a set of adverse consequences, such as low-grade inflammation, insulin resistance and oxidative stress, which in turn which affect grey and white matter integrity (Tchernof and Després, 2013). In this context, a previous longitudinal study examined BMI trajectories over 42 years and cortical thickness. Relative to normal-weight subjects, participants with a sustained history of obesity showed lower cortical thickness in the frontal pole, inferior frontal gyrus, inferior, middle and superior temporal gyri (Franz et al., 2019). Similarly, longitudinal increases in BMI have shown inverse correlations with cortical thickness in the posterior cingulate and supramarginal gyrus (Shaw et al., 2018), but also positive associations with volume of the hippocampus (Bobb et al., 2014). The possibility that longitudinal weight changes are associated with neuroanatomical variations in the reward system, however, remains largely unexplored.

Finally, a factor that could account for the low replicability of the findings is publication bias (Fusar-Poli et al., 2014). It has been suggested that findings implicating the ventral striatum might be overrepresented in the human neuroimaging literature (Behrens et al., 2013). Studies reporting significant effects in mesolimbic areas seem to have better chances of being published in peer-reviewed journals (Behrens et al., 2013; Ioannidis, 2011). At the same time, researchers might tend to underreport non-significant analyses (Ioannidis, 2011). These considerations warrant an objective overview of the relationship between NAcc volume and obesity and an examination of a possible publication bias.

Here, we first provide an evaluation of the available evidence reporting macroscopic changes in brain dopaminergic regions in obesity. Using a meta-analysis, we examined the consistency of results on obesity, possible publication bias and attempted to explain inconsistent findings in the literature. In the second part of the study, we examined whether BMI and BMI change over time are linked with differences in the NAcc and other reward-related regions. We addressed this last question by applying latent growth models in the UK Biobank dataset.

## Methods

### Meta-analysis

The literature search was conducted independently by IGG and FM, after which the results were cross-validated. We performed a literature search using Scopus database with two separate search terms: 1) obesity AND nucleus AND accumbens AND volume, and 2) obesity AND brain AND volume. Criteria for inclusion of studies were investigation of nucleus accumbens volume using MRI, inclusion of lean and obese individuals in the study, investigation of group differences in NAcc volume between lean and obese participants (grouped based on BMI) or correlations of NAcc volume and BMI or body fat percentage (BFP), and reporting of statistics necessary for further analysis. Whenever we detected sample overlap across studies, we included the study with the biggest sample size in the analysis. When studies reported separate results for left and right NAcc, we selected the association with the strongest effect size. To perform meta-analysis we used *Meta-essentials* software (Suurmond et al., 2017), which allows the use of results from regression and partial correlation analyses to calculate an overall effect size for all the studies together. From published studies investigating linear regressions/correlations of NAcc volume with BMI/BFP we collected the following statistics: partial correlation coefficients, regression beta values, standard errors of regression beta values, t-values, number of predictors in models, number of observations (sample size). For studies investigating group differences between lean and obese individuals we extracted t-values and converted them to correlation coefficients with the following equation: 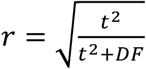, where *r* is the correlation coefficient and *DF* is the degrees of freedom. The obtained coefficients were then entered into the meta-analysis with a random effects model. We further investigated publication bias in the literature by means of a funnel plot and Egger test for funnel plot’s asymmetry (Egger et al., 1997; Egger and Smith, 1995). Lastly, due to the fact that included studies differed greatly with regard to the age range of participants, we performed a regression analysis between mean age and effect size (partial correlation) across studies.

### UK Biobank

We included 4924 participants from the UK Biobank. The UK Biobank (https://www.ukbiobank.ac.uk/) is a large-scale open access dataset recruited across UK. A subset of 25,000 participants to date have had extensive brain MRI measurements, health variables and clinical information, among other data (Miller et al., 2016). Participants’ ages range from 40 to 80 years and sex distribution is balanced. Participants in the imaging subset of the UK Biobank have undergone 3 visits: an initial assessment visit (2006-2010), a first repeat assessment visit (2012-2013) and a brain MRI imaging visit (started in 2014). In all the three visits participants provided information related to their general health, and researchers collected anthropometric measurements (i.e., body mass index or waist circumference).

We excluded subjects with psychiatric or neurological illness and cardiovascular problems (UK Biobank field IDs listed in the supplementary Table s1). Additionally, we excluded participants if their BMI was considered underweight (< 18.5 kg/m^2^, BMI field ID: 21001), since a very low BMI may be indicative of illness (Tchernof and Després, 2013).

To account for the possible effect of metabolic factors, we examined if participants met any of the following criteria considered to increase metabolic burden (Alberti et al., 2009): (a) high waist circumference (women: waist circumference >= 80; men: waist circumference >=94; calculated from ID: 48); (b) high blood pressure (systolic blood pressure >= 130 or diastolic blood pressure >= 85; extracted from ID: 4079/80); (c) past or current smoker (ID: 20116) and (d) diabetes (ID: 2443). For each of these criteria, participants scored 1 if they met the criterion and 0 if they did not. This way, we created a composite score that summed all the criteria for metabolic burden, ranging from 0 to 4 (Alberti et al., 2009; Yates et al., 2012). Other metabolic risk factors, such as cholesterolemia and high triglycerides, are not well characterized in the UK Biobank dataset and were not included. Similarly, we created a composite score to reflect affective symptoms. This score consisted of a sum of scores for the possible presence of (a) depression (ID: 4598); (b) anxiety (ID: 1980) and (c) anhedonia (ID: 4631). This score ranged from 0 to 3.

The inclusion/exclusion process is depicted in Figure 1. We conducted our research under the UK Biobank application number 35605.

**Figure 1.**
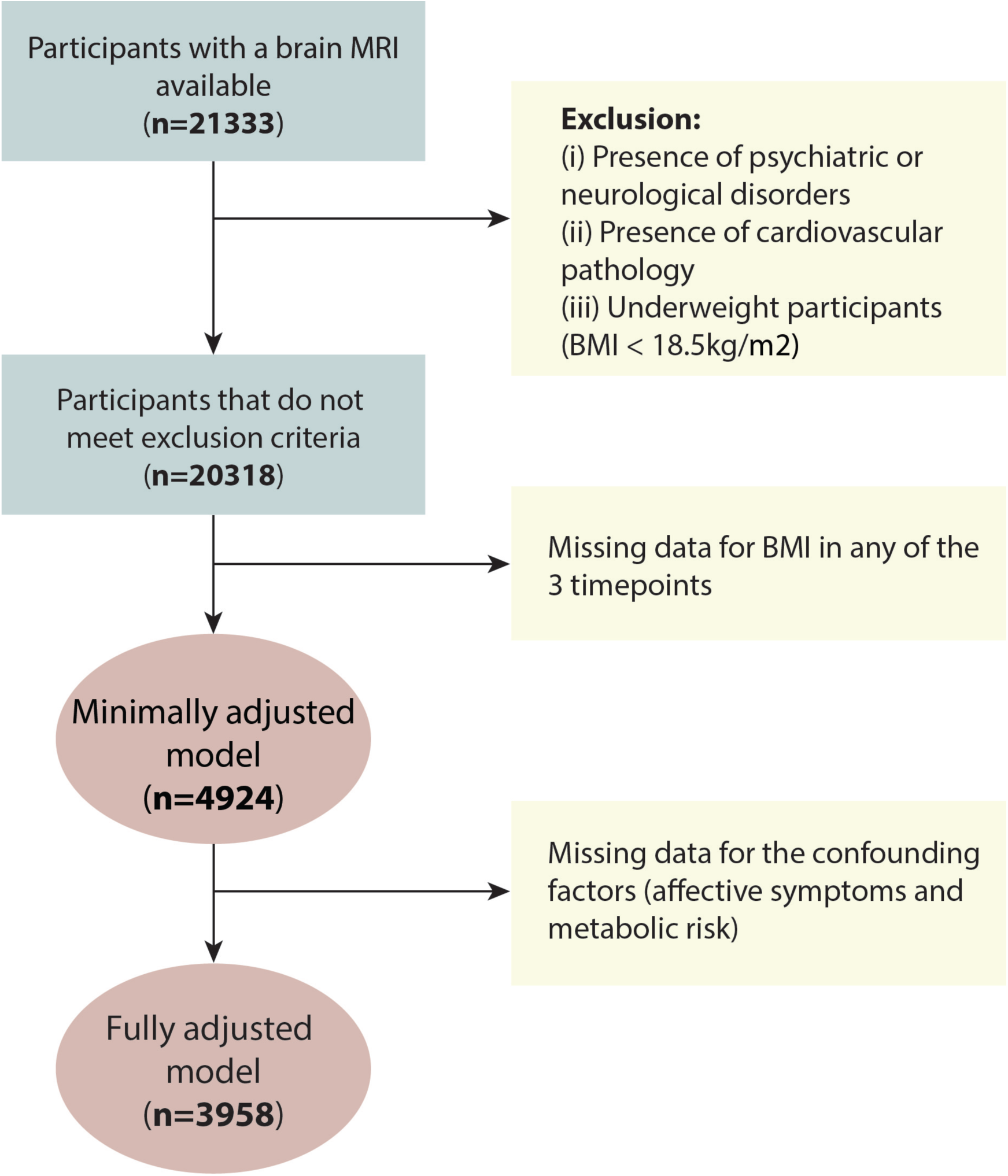
Description of the inclusion/exclusion process of the study.

### Regional brain volume

We used the brain volume variables available in UK Biobank. We selected as regions of interest (ROI) the NAcc (field ID: 25023/24) and other regions that are part of the mesolimbic circuitry: amygdala (ID: 25021 and 25022), caudate (ID: 25013 and 25014), hippocampus (ID: 25019 and 25020), pallidum (ID: 25017 and 25018), putamen (ID: 25015 and 25016), thalamus (ID: 25011 and 25012) and orbitofrontal cortex (ID: 25846 and 25847). For all regions, we added the volumes from the left and right hemisphere and divided them by total brain volume (field ID 25010).

### Statistical analysis: latent growth models in the UK Biobank dataset

Latent growth models can be used to describe how individuals differ in the longitudinal trajectory of variables of interest. In brief, latent growth models specify two variables: intercept and slope. The intercept depicts the initial point where each individual is located on the Y-axis. The slope represents their trajectory over time (Duncan and Duncan, 2009; Hertzog and Nesselroade, 2003).

Latent growth models have been applied to the UK Biobank dataset to examine longitudinal changes in fluid intelligence (Kievit et al., 2018). The model presented here is heavily inspired by this previous work.

We estimated the latent growth models using the Lavaan package in R (Rosseel, 2012). First, we built a basic model with the three available measurements of BMI. We estimated the latent variables’ intercept and slope from the repeated measurements of BMI. The intercept represents a constant across time for each individual, so the factor loading was fixed to 1. The slope was constrained to the mean intervals between time points (that is 0 for the first time point, 3.4 years for the second and 7.5 for the third). We included the brain areas defined as ROIs as regressors of the intercept and slope, and we allowed the ROIs to co-vary between each other and with age.

We assessed model fit with the chi-squared test, the root mean square error of approximation (RMSEA), the standardized root mean squared residuals (RMSR) and the comparative fit index (CFI). Goodness of fit was defined as: RMSEA (acceptable fit < 0.08, good fit < 0.05), CFI (acceptable fit 0.95 - 0.97, good fit > 0.97), SRMR (acceptable fit 0.05 - 0.10, good fit < 0.05) (Schermelleh-Engel et al., 2003).

To evaluate if simpler models would offer a better fit, we compared this basic model with two alternative models: (a) a model with the intercept constrained, which assumes that one average score can represent all the time-points, and (b) a model with the slope constrained, which assumes that individual scores do not change over time.

Finally, we built a fully adjusted model that controlled for the effects of metabolic risk and affective symptoms. We defined these two variables as regressors of the intercept and slope, and allowed them to co-vary between each other, with the brain ROIs and with age. We set the statistical threshold at p<0.005.

## Results

### Meta-analysis

We identified 17 samples within 14 studies eligible for the meta-analysis (Bernardes et al., 2018; Beyer et al., 2019; Coveleskie et al., 2015; Curran et al., 2013; de Groot et al., 2017; Dekkers et al., 2019; Hamer and Batty, 2019; Annette Horstmann et al., 2011; Kakoschke et al., 2019; Mole et al., 2016; Ottino-González et al., 2017; Perlaki et al., 2018; Rapuano et al., 2017; Widya et al., 2011). For detailed description of the studies see Table 1. Two studies used samples from the UK Biobank dataset (Dekkers et al., 2019; Hamer and Batty, 2019), hence we only included the study with the larger sample size in the meta-analysis (Dekkers et al., 2019). Two of the studies presented separate results for men and women, which were introduced into the meta-analysis as four separate studies (Dekkers et al., 2019; Annette Horstmann et al., 2011). A study by Kakoschke et al., 2019, included adolescent and adult samples, which we also included as two separate studies in the meta-analysis.

**Table 1.**
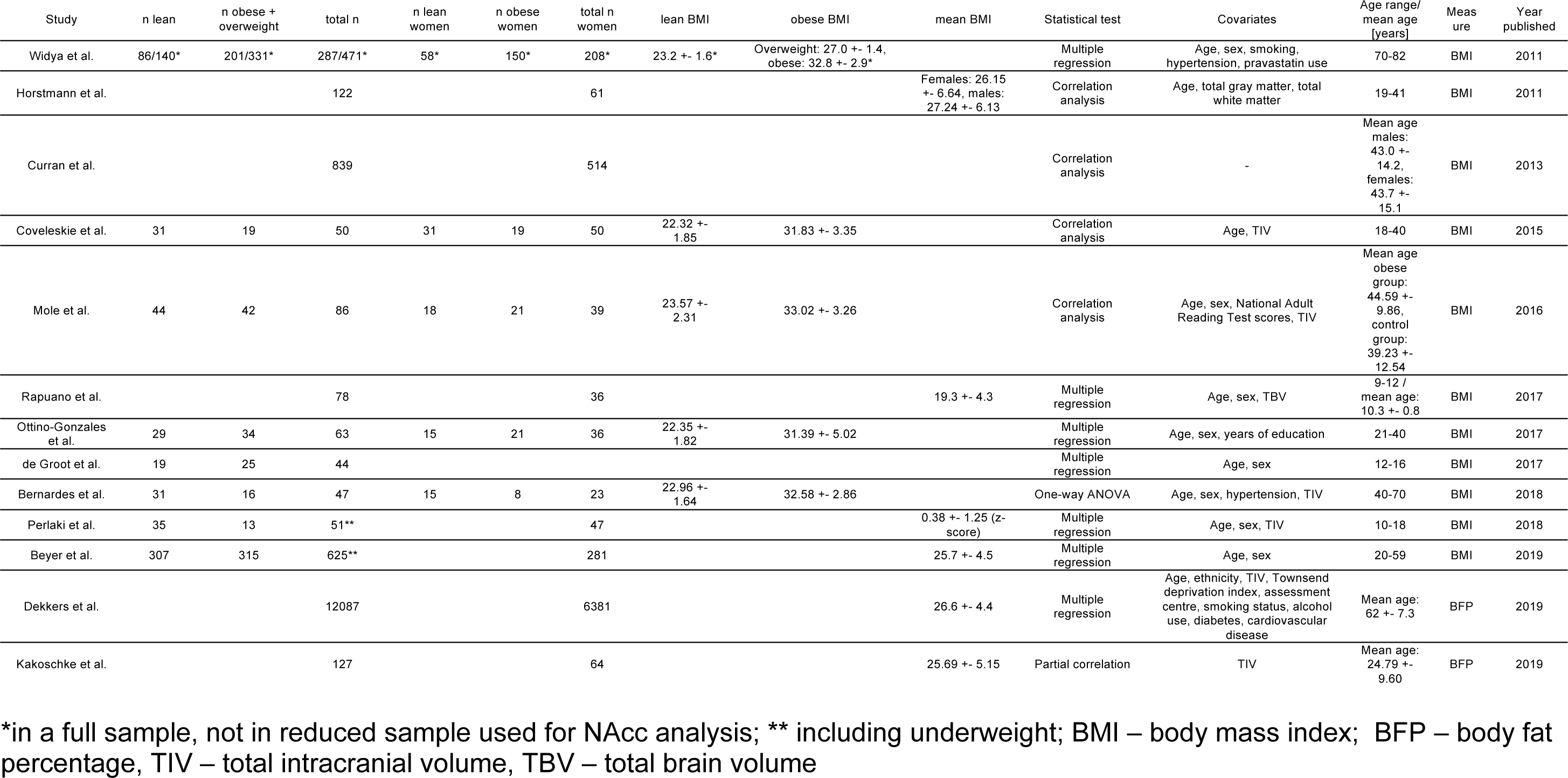
Details of studies included in the meta-analysis

The meta-analysis points to a significant positive association between BMI and NAcc volume (mean partial correlation value r=0.15, confidence intervals: 0.04-0.27, Fig. 2). Heterogeneity values in the investigated studies was I^2^=91.50%. Funnel plot and Egger test did not provide evidence for publication bias (t=1.12, p=0.283; Fig. 3). Investigation of the relationship between partial correlation values and age of participants for each study pointed to a significant association between the two variables (β=-0.60, p<0.001, Fig. 4). Here, lower age was related to a positive correlation between BMI and NAcc volume, while higher age showed an opposite pattern.

**Fig. 2.**
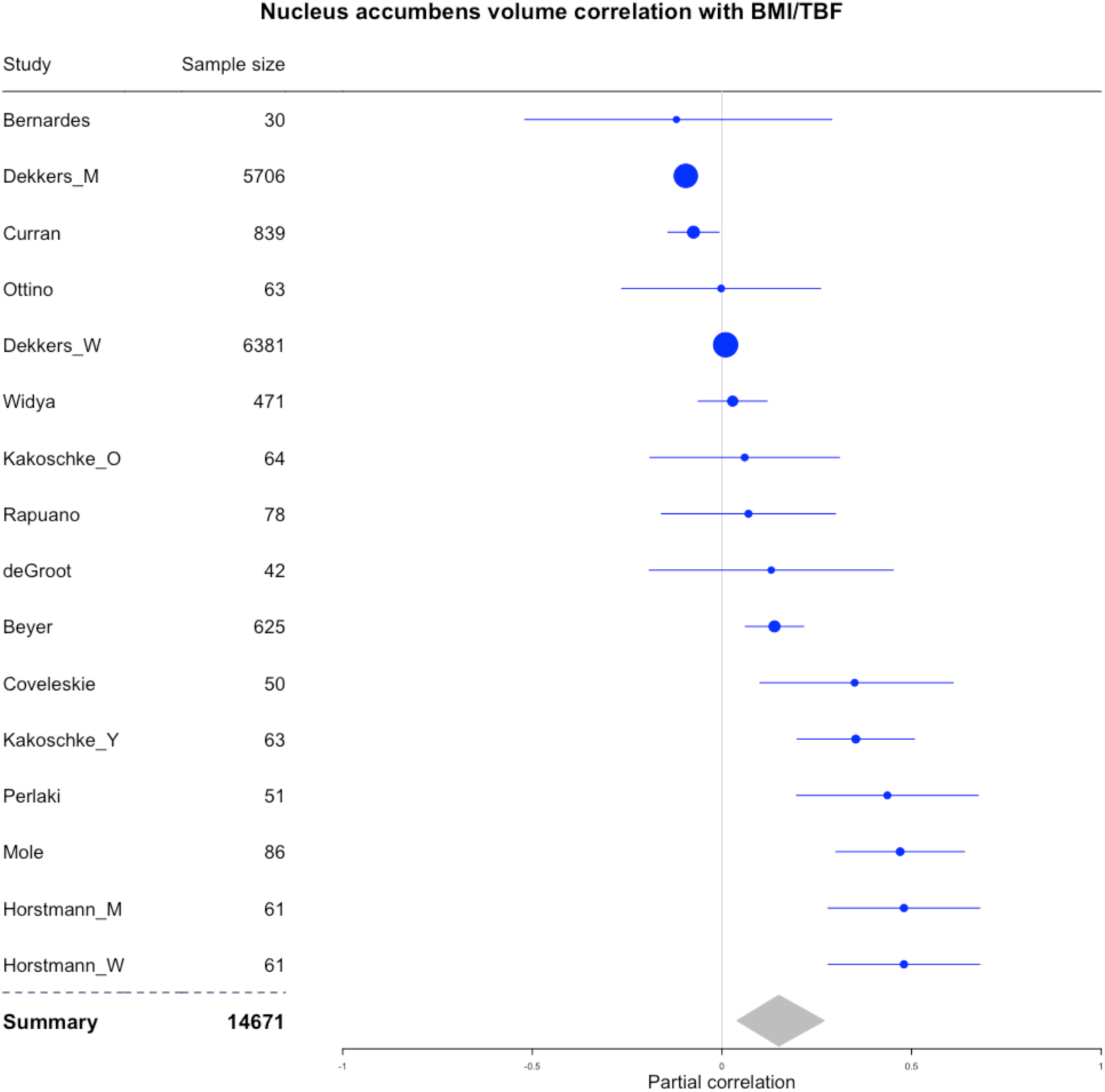
Correlation coefficients and 95% confidence intervals for relationships between BMI/BFP and nucleus accumbens volume for each study included in the meta-analysis. Sizes of points are proportional to sample sizes. Summary diamond represents overall partial correlation for all studies. M – men, W – women, Y – adolescent sample, O – adult sample.

**Fig. 3.**
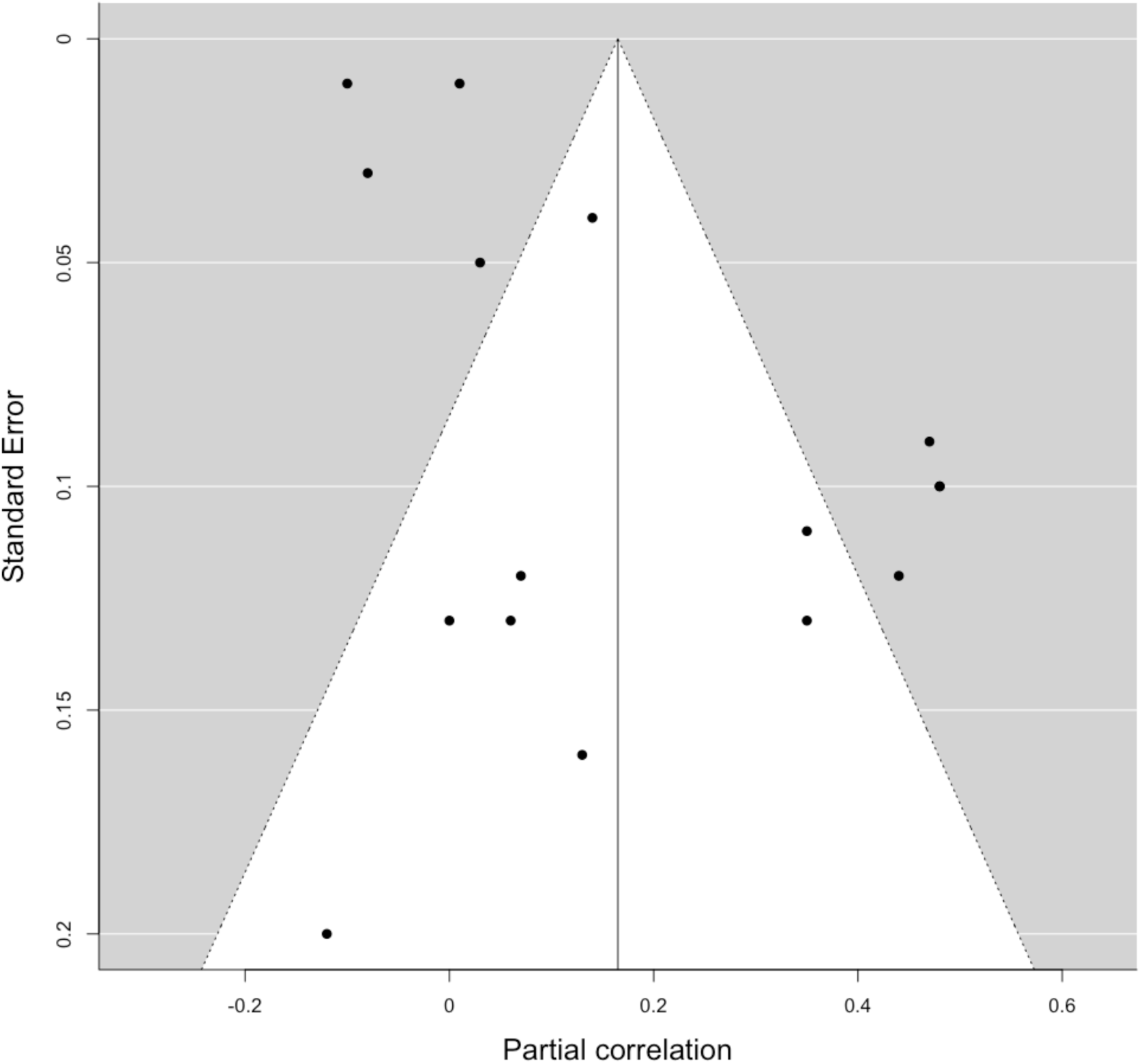
Publication bias funnel plot. Dots represent each study included in the meta-analysis. Note that there are only 15 dots, while we included 16 separate results (from 13 studies); this is due to the fact that in Horstmann et al. results for men and women were the same, so one of the dots represents two studies.

**Fig. 4.**
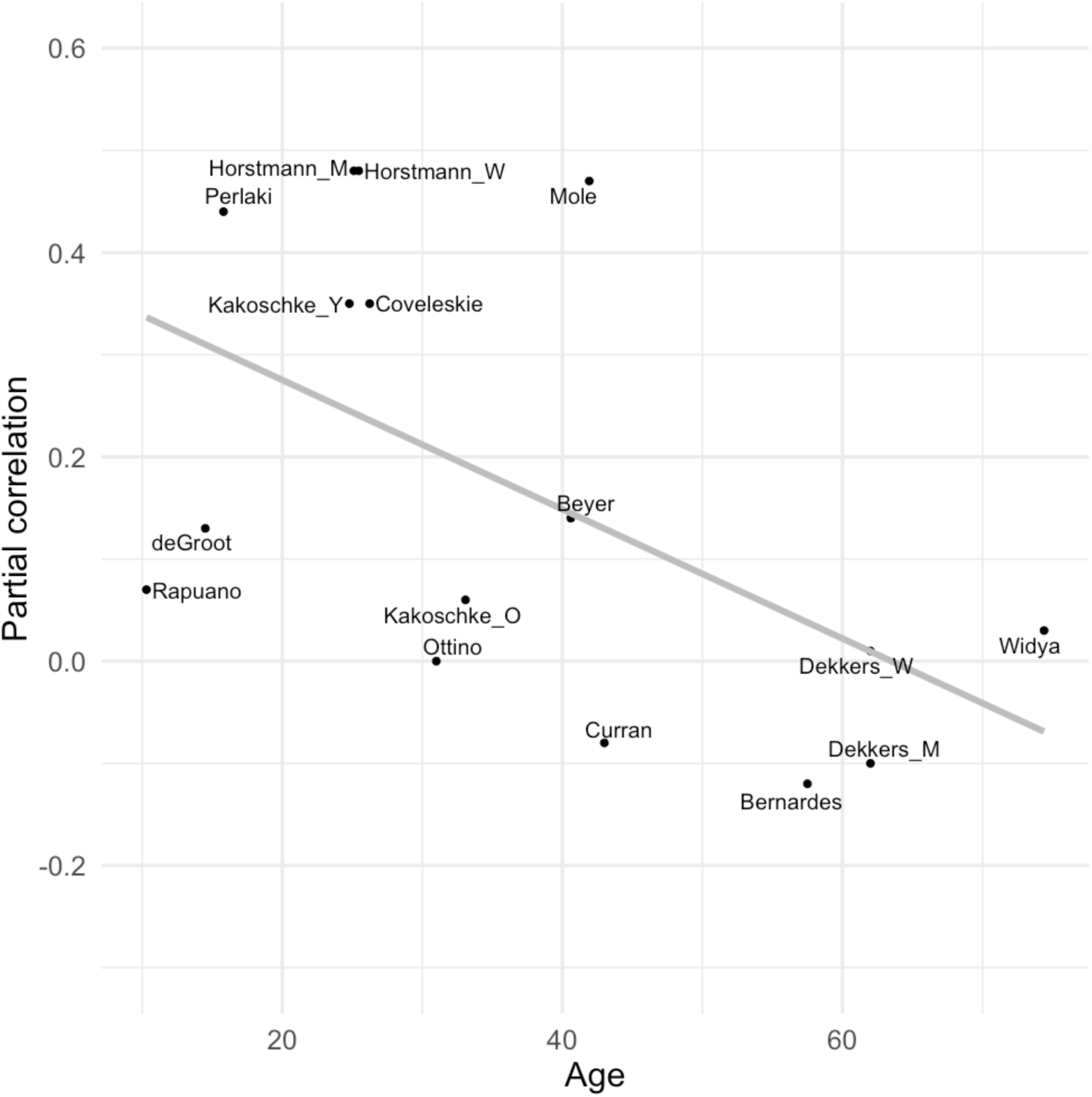
Relationship between partial correlation and age in all studies included in the meta-analysis. Each study is represented by a dot. Grey line represents the best fit. M – men, W – women, Y – adolescent sample, O – adult sample

### Latent growth model in the UK Biobank dataset

Table 2 presents an overview of the demographic and anthropometric characteristics of the participants.

**Table 2.** Demographic and anthropometric characteristics of the participants included in UK Biobank latent growth analysis (2593 women and 2394 men).

|  | Minimum | Maximum | Mean | Standard deviation |
| --- | --- | --- | --- | --- |
| Age at timepoint 1 | 40 | 70 | 55.62 | 7.37 |
| Age at timepoint 2 | 44 | 74 | 59.92 | 7.31 |
| Age at timepoint 3 | 46 | 80 | 63.24 | 7.38 |
| Body Mass Index at timepoint 1 | 18.63 | 55.50 | 26.51 | 4.20 |
| Body Mass Index at timepoint 2 | 18.58 | 54.74 | 26.55 | 4.21 |
| Body Mass Index at timepoint 3 | 18.52 | 53.36 | 26.37 | 4.24 |
|  | **Percentage** |  |  |  |
| Sum of affective symptoms (depression, anxiety and anhedonia) | Zero: 29.0%<br>One: 31.6%<br>Two: 21.8%<br>Three: 17.6% |  |  |  |
| Sum of metabolic symptoms (high waist circumference, hypertension, diabetes, current or past smoker) | Zero: 21.7%<br>One: 43.8%<br>Two: 31.1%<br>Three: 3.4%<br>Four: 0.0% |  |  |  |

#### Basic model: Longitudinal changes in BMI

Our minimally adjusted model depicted the association between longitudinal changes in BMI and the mesolimbic regions of interest. We included as covariates age and our brain regions of interest. The model provided a good fit (χ^2^(10)=56.14; RMSEA=0.031; SRMR=0.003; CFI=0.998).

The mean and variance of the intercept were 36.44 and 16.50, respectively. The mean slope showed a non-significant positive trend (estimate=0.22; SE=0.09; p=0.016) suggesting that participants’ weight remained relatively stable over time. The slope variance was positive (estimate=0.040; SE=0.006; beta=0.990; p<0.001). There was a negative effect of age on slope (estimate=-0.004; SE=0.001; beta=-0.099; p<0.001), suggesting the existence of age differences in the slope. The intercept and the slope showed a negative covariance (estimate=-0.113; SE=0.027; beta=-0.15; p<0.001), suggesting that participants with a higher baseline BMI gained less weight. The intercept of BMI was significantly associated with the NAcc (estimate=-0.19; SE=0.04; beta=-0.08; p<0.001) and the thalamus (estimate=-0.49; SE=0.01; beta=-0.09; p<0.001). No areas showed a significant association with the slope (Figure 5 A).

**Figure 5.**
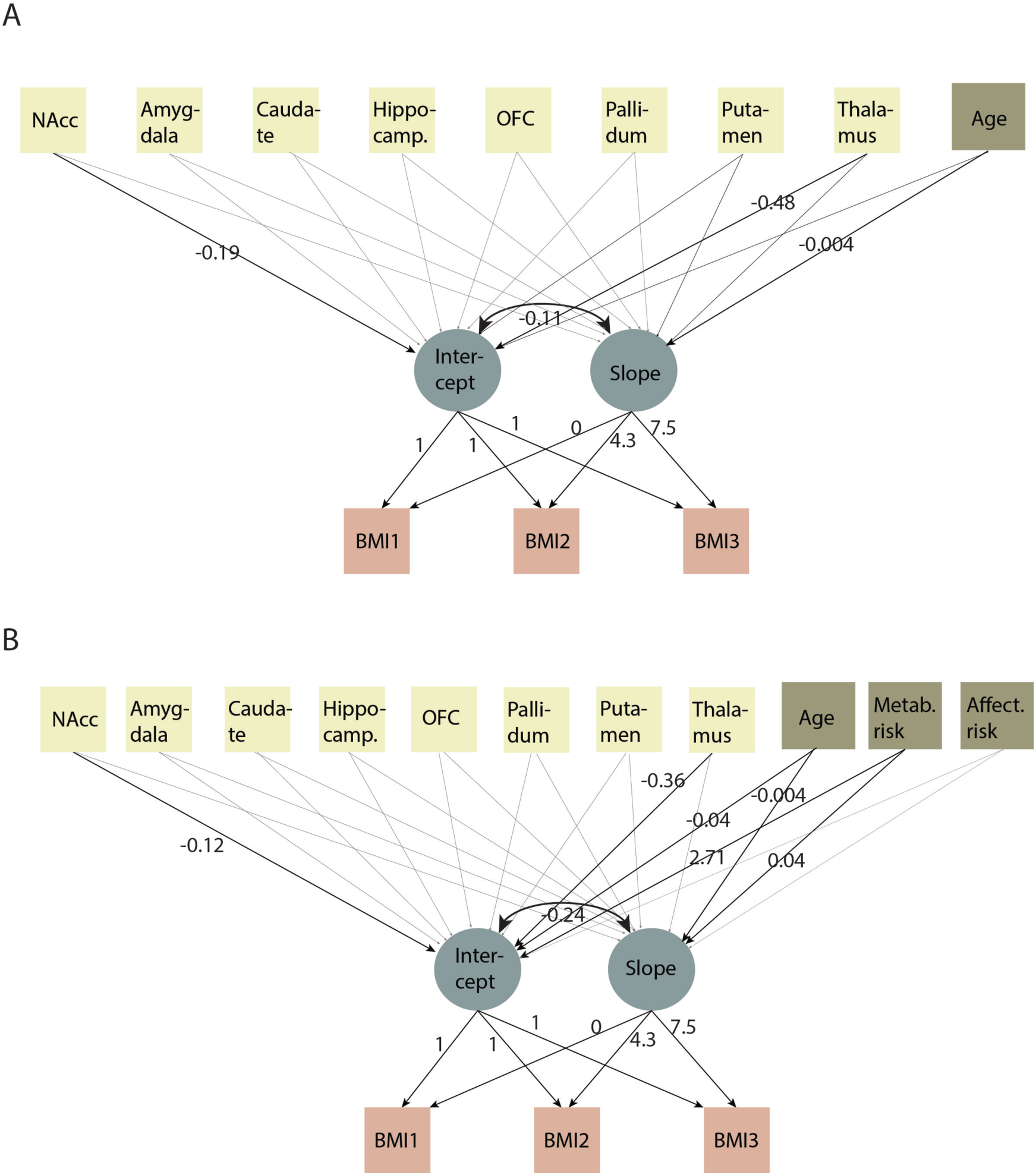
Latent growth models for BMI change across the 3 timepoints. A) Minimally-adjusted model; B) Fully-adjusted model. The values represent the estimate. Brain regions, age, metabolic risk and affective risk were allowed to co-vary between them (for simplicity, these associations are not represented).

#### Alternative (simpler) models

There was a significant decrease in model fit both when constraining the intercept (χ^2^(11) = 5351.2, p=2.2e-16) and when constraining the slope (χ^2^(11) = 53.926; p=2.08e-13), indicating that these alternative models did not provide a better solution.

#### Fully adjusted model (addition of metabolic risk and affective symptoms in the model)

We built a fully adjusted model by adding metabolic risk score and affective symptoms score as covariates. The model provided a good fit (χ^2^(12)=54.96; RMSEA=0.030; SRMR=0.003; CFI=0.998). The associations between NAcc and thalamus and the intercept remained equivalent (NAcc: estimate= −0.12; SE=0.04; beta=-0.05; p=0.004; thalamus: estimate=-0.37; SE=0.09; beta=-0.07; p<0.001) (Figure 5 B).

#### Examination of results by sex

To explore the possibility that sex differences might be driving the results, we tested the fully adjusted model dividing the sample in men and women. The models produced equivalent results in terms of effect size compared to the model that aggregates both sexes (data not shown for simplicity).

## Discussion

In the present study we sought to determine whether obesity is associated with volumetric differences in the NAcc and other mesolimbic areas. To do so, we performed a meta-analysis on 15 independent samples, representing 14671 participants with available measurements of adiposity (i.e., BMI or total body fat). We found that, overall, NAcc volume was positively related to obesity measures. The magnitude of the correlation was small (r=0.16). There was no evidence of publication bias across the studies, according to the funnel plot and Egger test. We additionally examined possible age effects on the meta-analytic results, since the studies had a broad age range (9 to 82 years old). We found that the relationship was strongly dependent on age. Here, young participants showed a positive relationship between NAcc volume and obesity measures, while in older adults the relationship was negative. Next, using the UK Biobank dataset we tested whether longitudinal change in BMI was related to volumetric differences in the NAcc and in other mesolimbic regions. We included 4924 participants with available neuroimaging data that completed an evaluation of BMI across three time-points, covering 7.5 years. We applied latent growth models and estimated two latent variables, representing individual variations in BMI (i.e., intercept) and BMI change over time (i.e., slope). Contrary to our expectations, none of the regions of interest showed associations with BMI change over time.

However, the NAcc and thalamus showed negative associations with BMI. Our findings align well with our meta-analytic results, since the age range of the participants in the UK Biobank is 40 to 80 years old, and replicate previous results from this dataset (Dekkers et al., 2019).

The meta-analytic findings regarding an effect of age on the relationship between BMI/BFP and nucleus accumbens volume might help explain conflicting findings in the literature. First, increased NAcc volume in children and young adults with higher BMI might constitute a vulnerability factor for obesity. In fact, a study by Rapuano and colleagues (2017) showed that children with a higher risk for obesity as defined by an FTO gene polymorphism have greater fMRI responses to viewing food commercials in the NAcc (Rapuano et al., 2017). Additionally, in the same study, children at a higher risk for obesity showed larger NAcc volumes. This is in line with studies in children and adolescents showing positive relationships between BMI/BFP and NAcc volume (de Groot et al., 2017; Perlaki et al., 2018).

On the other hand, a negative association between BMI/BFP and NAcc volume in older adults might be related to age-related increases in vascular pathology and inflammatory processes. Previous animal studies showed that diet-induced obesity influences structure and function of the NAcc (Gutiérrez-Martos et al., 2018), but also other brain areas, through neuroinflammation (Beilharz et al., 2016; Lu et al., 2011; Tapia-González et al., 2011). Our findings are thus in line with a study in humans showing that total brain volume is negatively associated with inflammatory markers (Jefferson et al., 2007) and a study showing that, in older adults, high BMI is generally related to brain atrophy (Raji et al., 2009).

Such an age-dependent relationship between NAcc volume and BMI might act as a confound in nuclear imaging studies on dopamine receptor density in obesity, due to partial volume effects: decreased NAcc volume in older adults with higher BMI might cause increased measurement error. It has been suggested that dopaminergic dysfunctions in mesolimbic pathways facilitate weight gain and obesity through alterations in reinforcement learning, motivation and self-regulation (Volkow et al., 2017). However, results in obesity are highly heterogeneous, with studies reporting reductions (Wang et al., 2001), null findings (Eisenstein et al., 2015) and increases (Gaiser et al., 2016) in striatal D2/D3 receptor availability in obesity. A study by Dang and colleagues reported null associations between striatal D2/D3 receptor availability and BMI in individuals younger than 30 years old and positive associations in the midbrain, putamen and ventral striatum in participants older than 30 (Dang et al., 2016). It is possible that nucleus accumbens volume differences between age groups might bias D2/D3 receptors measurements. Hence, those differences might be important in explaining the age-dependent relationship between BMI and D2/D3 receptor availability found by Dang and colleagues.

A history of consistently elevated adiposity increases morbidity and mortality. Specifically, as the number of years lived with obesity increases, there is higher incidence of all-cause, cardiovascular-related and cancer-related mortality (Abdullah et al., 2011b). The incidence of Type 2 diabetes also increases by 12% to 14% for each 2 years of increment in the duration of obesity (Abdullah et al., 2011a; Hu et al., 2014). In this context, we further examined the possible association between longitudinal changes in BMI and higher brain vulnerability to the effects of adiposity. For this analysis, we used the UK Biobank cohort, which includes participants with an age range of 40 to 80 years. In this well-powered sample, we did not find evidence that longitudinal changes in BMI are associated with mesolimbic volumetric differences. However, we found that the intercept of BMI (which represents mean BMI) was negatively associated with the volume of the NAcc and thalamus. These associations remained significant when accounting for vascular risk factors and affective symptoms in the analysis. This indicates that a high BMI is associated with lower NAcc and thalamus volume in older adults, possibly reflecting detrimental effects of adiposity on the brain. The main limitation of our meta-analysis is the fact that it includes only cross-sectional measures. Ideally, to be able to confirm our interpretation of the results, a longitudinal study investigating the relationship between BMI/BFP and NAcc volume across the lifespan should be performed. Such studies, even though difficult to perform, become more feasible with the development of large cohorts. Although we hypothesized that inflammatory markers might mediate the relationship between obesity and brain vulnerability, we could not test this hypothesis since our data did not contain plasma inflammatory markers. We hope that future neuroimaging studies will address this issue.

## Conclusions

In the present study we have examined the relationship between obesity measurements and volumetric differences in the NAcc. Using meta-analysis, we found that obesity showed a small positive relationship with gray matter volume in the NAcc. We did not find evidence of publication bias. Moreover, a further analysis indicated that age was a moderator of this association. In younger participants the associations between obesity and NAcc volume are positive, while in older participants this relationship is null or negative. Next, in a population-based cohort of older adults we tested whether longitudinal changes in BMI are related to volumetric differences in the NAcc and other mesolimbic areas. Contrary to our expectations, none of the areas examined showed associations with change in BMI over time. However, the NAcc and the thalamus volumes showed negative correlations with individual BMI, supporting the meta-analysis results. We point to the possible implication of inflammatory factors, which might be especially relevant in individuals in their mid and older adulthood.

## Acknowledgements

We thank Frauke Beyer, Jonatan Ottino-González and Kristina Rapuano for providing us with additional information from their studies to complete the meta-analysis. This work was supported by a Foundation Scheme award to AD from the Canadian Institutes of Health Research. IGG is a recipient of a Postdoctoral Fellowship from the Canadian Institutes of Health Research.

