## supplementary Table s1 for "Nucleus accumbens volume is related to obesity measures in an age-dependent fashion"

| **Supplementary Table 1.** ID fields from the UK Biobank used for exclusion criteria | | | |
| --- | --- | --- | --- |
| **Health condition** |  | **Field** | **Subfield or code** |
| Bipolar disorder status |  | 20122 | "Bipolar Type I (Mania)" or "Bipolar Type II (Hypomania)" |
| Other mental health problems | | 20544 |  |
|  | Personality disorder |  | "A personality disorder" |
|  | Schizophrenia |  | "Schizophrenia" |
| Vascular heart diagnoses |  | 6150 | "Heart attack", "Angina" or "Stroke" |
| Neurological disorders |  | 20002 |  |
|  | Dementia or Alzheimer’s disease | | 1263 |
|  | Parkinson’s disease |  | 1262 |
|  | Chronic degenerative neurological | | 1258 |
|  | Guillain-Barré syndrome |  | 1256 |
|  | Multiple Sclerosis |  | 1261 |
|  | Other demyelinating disease | | 1397 |
|  | Stroke or ischaemic stroke | | 1081 |
|  | Brain cancer |  | 1032 |
|  | Brain haemorrhage |  | 1491 |
|  | Brain/intracranial abscess | | 1245 |
|  | Cerebral aneurysm |  | 1425 |
|  | Cerebral palsy |  | 1433 |
|  | Encephalitis |  | 1246 |
|  | Epilepsy |  | 1264 |
|  | Head injury |  | 1266 |
|  | Infections of the nervous system | | 1244 |
|  | Ischaemic stroke |  | 1583 |
|  | Meningeal cancer |  | 1031 |
|  | Meningioma (benign) |  | 1659 |
|  | Meningitis |  | 1247 |
|  | Motor Neuron Disease |  | 1259 |
|  | Neurological injury/trauma | | 1240 |
|  | Spina bifida |  | 1524 |
|  | Subdural haematoma |  | 1083 |
|  | Subarachnoid haemorrhage | | 1086 |
|  | Transient ischaemic attack | | 1082 |
